# Vulnerability and adaptations to climate change in EU protected areas: a Natura 2000 managers’ perspective

**DOI:** 10.64898/2025.12.19.695111

**Authors:** Giorgio Zavattoni, Jon E. Brommer, Tyler Hallman, Ineta Kačergytė, Szabolcs Nagy, Tomas Pärt, Diego Pavón-Jordán, Alaaeldin Soultan, Yanjie Xu, Elie Gaget

## Abstract

The Natura 2000 (N2K) network is a key conservation instrument in the European Union, but its effectiveness is challenged by climate change. We surveyed 382 N2K managers to investigate their perceptions of climate change and related site adaptation strategies. Warming and precipitation shifts were frequently perceived as threats, with vulnerability of N2K sites highest in the Mediterranean and lowest in the Boreal region. Our results suggest that the official N2K site information stored in the Standard Data Forms greatly under-report managers’ assessment of vulnerability. Around half of the surveyed managers implemented adaptation strategies, which, when characterised following a Resist-Accept-Direct framework, aimed not only at resisting, but also at directing and accepting the effects of climate change. Managers also highlighted the need for scientific knowledge on local vulnerability and increased funding to improve the implementation of adaptation strategies inside protected areas.

## Introduction

Anthropogenic climate change is a growing threat to biodiversity (IPBES 2019). Protected areas can help species cope with this change by maintaining healthy ecosystems (Bowgen et al. 2022; Gaget et al. 2021), though climate change mitigation is rarely a primary target of their management. With the intensifying of climate change, conservation efforts need to explicitly align with its effects to avoid failures and waste of limited conservation resources (Greenwood et al. 2016).

To help management decisions under climate change, the Resist-Accept-Direct (RAD) framework has been theorized (Schuurman et al. 2020). The RAD framework specifies that climate change adaptation strategies can aim to: 1) resist the effects of climate change, by maintaining locally historical ecosystem condition; 2) accept the effects, by not influencing climate-imposed changes; or 3) direct the effects, by guiding the changes towards an alternative and desirable condition. For example, if a forest is degrading due to warming, this could be resisted by planting local tree species, accepted by allowing the degradation, or directed by translocating tree species adapted to warmer temperatures (Magness et al. 2022). Choosing the right strategy depends on conservation goals, and the three options can be implemented at the same site for different purposes. The Natura 2000 (N2K) is an extensive network of protected areas across the European Union (EU), and each site has clearly defined conservation targets (Kati et al. 2015). However, there is uncertainty about the effectiveness of this network in the face of climate change (Araújo et al. 2011), with many N2K sites projected to experience new climatic conditions (Nila et al. 2019). A study on forest ecosystems in Belgium found that around one third of N2K managers accounted for climate change in their work (Sousa-Silva et al. 2016) while in France, the Natur’Adapt project (LIFE17 CCA/FR/000089), found that only 15% of the managers did so (EC 2025).

However, to our knowledge, no large-scale studies investigated how managers perceive biodiversity vulnerability to climate change across Europe, which climatic drivers they feel threaten biodiversity (e.g., warming, precipitation shifts), how they adapt management practices accordingly, and what challenges they face in doing so. To fill this gap, we conducted a pan-European survey targeting N2K managers, asking about their perception of site vulnerability to climate change-related threats, the implemented adaptation strategies, and the barriers they face in implementing them. We investigate how often different aspects of climate change are perceived as threats to biodiversity, and we compare these data to the latest official information reported to the European Commission (EC) for each N2K site through the Standard Data Forms (SDF). We hypothesize that vulnerability to climate change differs across biogeographical regions and that vulnerable N2K sites will be more likely to implement adaptation strategies. Following the RAD framework, we expect that implemented climate change adaptation strategies have primarily aimed to resist the effects of climate change due to the traditional static approach to conservation. Our results provide insights on implemented strategies for climate change adaptations at the EU scale and help to identify barriers for their applications.

## Methods

### The survey

We built an online survey to gather information on how N2K managers perceive and address climate change (see Appendix S1 and S2 for details). Managers selected an N2K site under their supervision and answered different questions (Table 1; Appendix S2). Multiple answers from the same site (21 instances) were considered separately.

**Table 1.** Questions of the survey section containing climate-change related topics.

| Category | Question | Possible answers |
| --- | --- | --- |
| **1. Vulnerability** | Do you think the increase in temperature is a threat for species/habitats targeted in this protected area? | "Yes", "No", "Probably in the future" |
|  | Do you think the change in the frequency of precipitations is a threat for species/habitats targeted in this protected area? | "Yes", "No", "Probably in the future" |
|  | Do you think sea level rise is a threat for species/habitats targeted in this site? | "Yes", "No", "Probably in the future" |
|  | Do you think the increased frequency of climatic extreme events is a threat for species/habitats targeted in this protected area? | "Yes", "No", "Probably in the future" |
| **2. Adaptations** | Do you consider the impact of climate change in the management of the Natura 2000 site? | "Yes", "No", "I do not know" |
| **3. RAD**<br>(Considered only if the answer to 2. was "Yes"; preceded by a brief explanation of the RAD framework) | Which action(s), if any, have been done in the site to RESIST the effects of climate change? | List of previously selected actions that are being implemented at the site |
|  | Which action(s), if any, have been done in the site to ACCEPT the effects of climate change? | List of previously selected actions that are being implemented at the site |
|  | Which action(s), if any, have been done in the site to DIRECT the effects of climate change? | List of previously selected actions that are being implemented at the site |
| **4. Implementation barriers**<br>(Considered only if the answer to 2. was "Yes") | Is there something preventing you from better implementing actions to deal with the effects of climate change on your site? | "No", "Knowledge on site vulnerability", "Knowledge on conservation actions useful to implement the RAD framework", "Capacity building", "Funding", "Time", "Consensus with stakeholders", "Other" |

We downloaded biogeographical region borders (EEA 2025) to investigate if respondents’ answers spatially differed. Sites under multiple regions were assigned to the terrestrial region with the highest coverage. One site located within the Marine Macaronesian region was excluded. Three sites under two regions without reported coverage were considered for both. The Steppic and Black Sea regions were merged due to their small size and proximity.

We downloaded the latest official site-level threat information (EEA 2021) to compare the vulnerability reported by the managers answering the survey with what was officially reported in the SDFs. One site was considered vulnerable to climate change if at least one of the four investigated climate aspects (warming, precipitation changes, extreme events, sea level rise) was reported as a threat.

### Statistical analysis

We used a generalized linear mixed model (GLMM) to test the probability of a site being perceived as vulnerable to the four aspects of climate change (warming, precipitation changes, extreme events, sea level rise). We used the type of climate change aspect as predictor and their reported vulnerability as binary response. Since every N2K site had one row for each of the four climate change aspects, the site was included as random effect.

We tested for biogeographical region variations on the vulnerability to warming, precipitation changes, extreme events, and overall vulnerability to climate change (i. e., vulnerability to at least one of the three aspects) by running a generalized linear model (GLM, binomial) for each of them (4 models), with biogeographical regions as predictors. We excluded sea level rise due to uneven coastal sites distributions between regions and answers from the Pannonian region (11) due to model convergence issues.

To assess if perceived vulnerability to climate change motivated the implementation of adaptation strategies, we fitted three binomial GLMs. In each GLM, overall vulnerability was the predictor and the three responses were the answers to the question about adaptation strategies: 1) implemented (yes/no); 2) not implemented (yes/no); 3) implementation uncertain (yes/no).

To examine differences in the implementation of RAD approaches, we fitted a binomial GLMM. We used the approach type (resist, accept, or direct) as predictor and their implementation (yes/no) as response. Since each N2K site was repeated 3 times (one for each approach), the site was included as random effect. Similarly, we tested if the frequency of implementation barriers differed. These two analyses were restricted to sites where adaptation strategies were reported.

All analyses were performed using R.4.2.1 (R Core Team 2022). The models were fitted with the ‘sdmTMB’ package correcting for spatial autocorrelation through a Gaussian random field with a cutoff distance of 0.1 decimal degrees (Anderson et al. 2024). The absence of spatial autocorrelation was validated with a Moran’s I test with threshold value of *p* = 0.05 (‘DHARMa’ package; Hartig 2022). P-values were calculated based on a Welsh test and pairwise comparisons extracted with the ‘emmeans’ package (Lenth 2023).

## Results

We obtained 382 answers from 25 different countries (see Zavattoni et al. 2025 for survey details).

### Vulnerability to climate change

Vulnerability to climate change was reported in 59.7% of the answers (228/382). Vulnerability to precipitation changes and warming were the most reported, followed by extreme events and sea level rise (Figure 1; Appendix S3).

**Figure 1.**
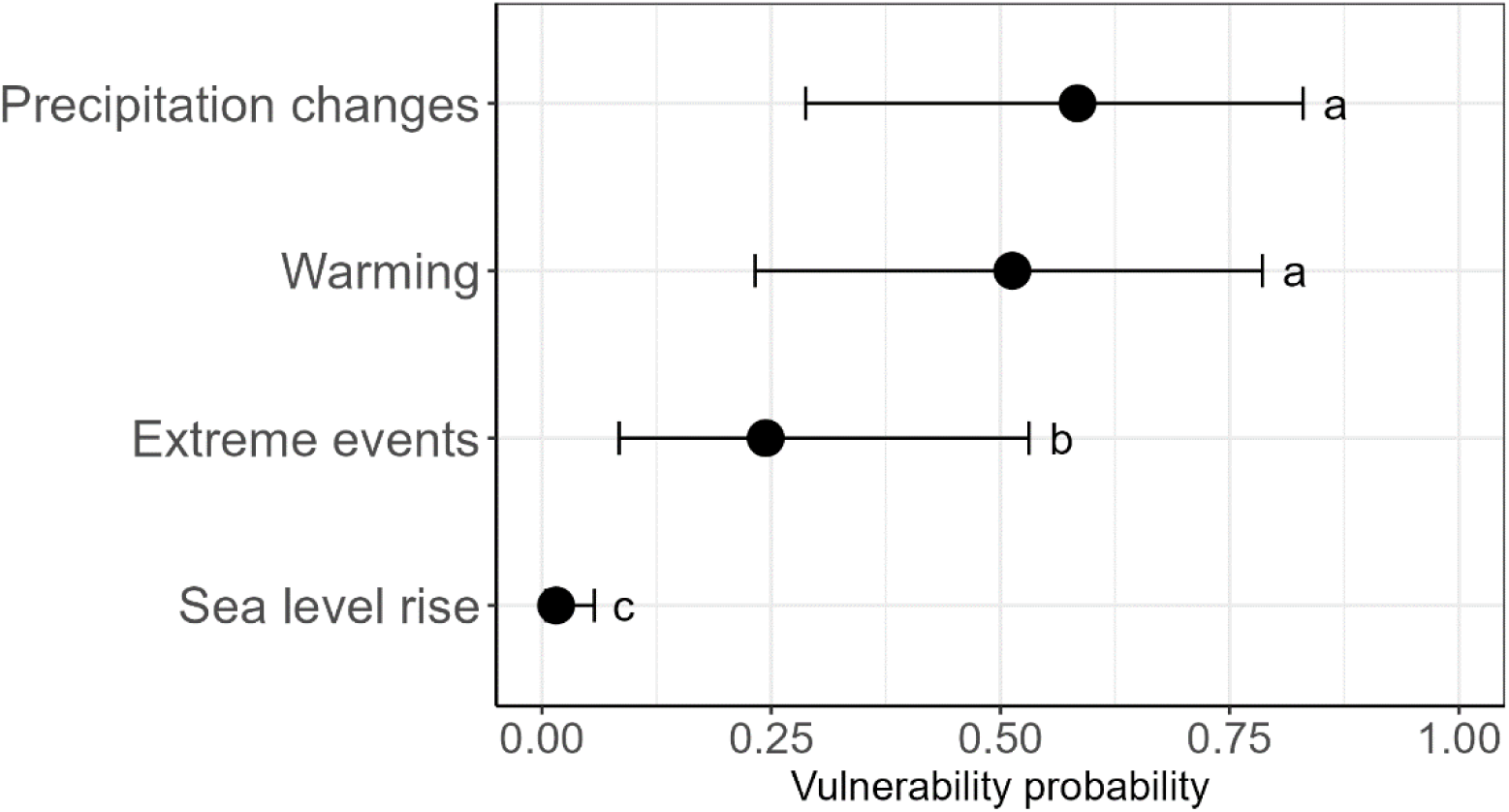
Predicted probabilities (± 95% CI) that a site reported vulnerability to different aspects of climate change. Vulnerability categories whose probabilities are statistically different from each other at *p* < 0.05 are marked with different letters, i.e., shared letters indicate no difference.

Among responses reporting vulnerability to climate change, 7.0% (16/228) came from N2K sites whose SDFs mentioned climate change as a threat, while 4.5% (7/154) of responses that did not report it as a threat, came from sites whose SDFs mentioned it.

Overall perceived vulnerability to climate change was not significantly different across biogeographical regions (Figure 2b; Appendix S4). Respondents from the Mediterranean region perceived higher vulnerability to warming (*p* = 0.02; Figure 2b; Appendix S5) and to precipitation changes (*p* = 0.03; Figure 2b; Appendix S6) than respondents from the Boreal region. Both Boreal and Continental regions were perceived as less vulnerable to extreme events than the Mediterranean (*p* < 0.01 and *p* = 0.03, respectively; Figure 2b; Appendix S7).

**Figure 2.**
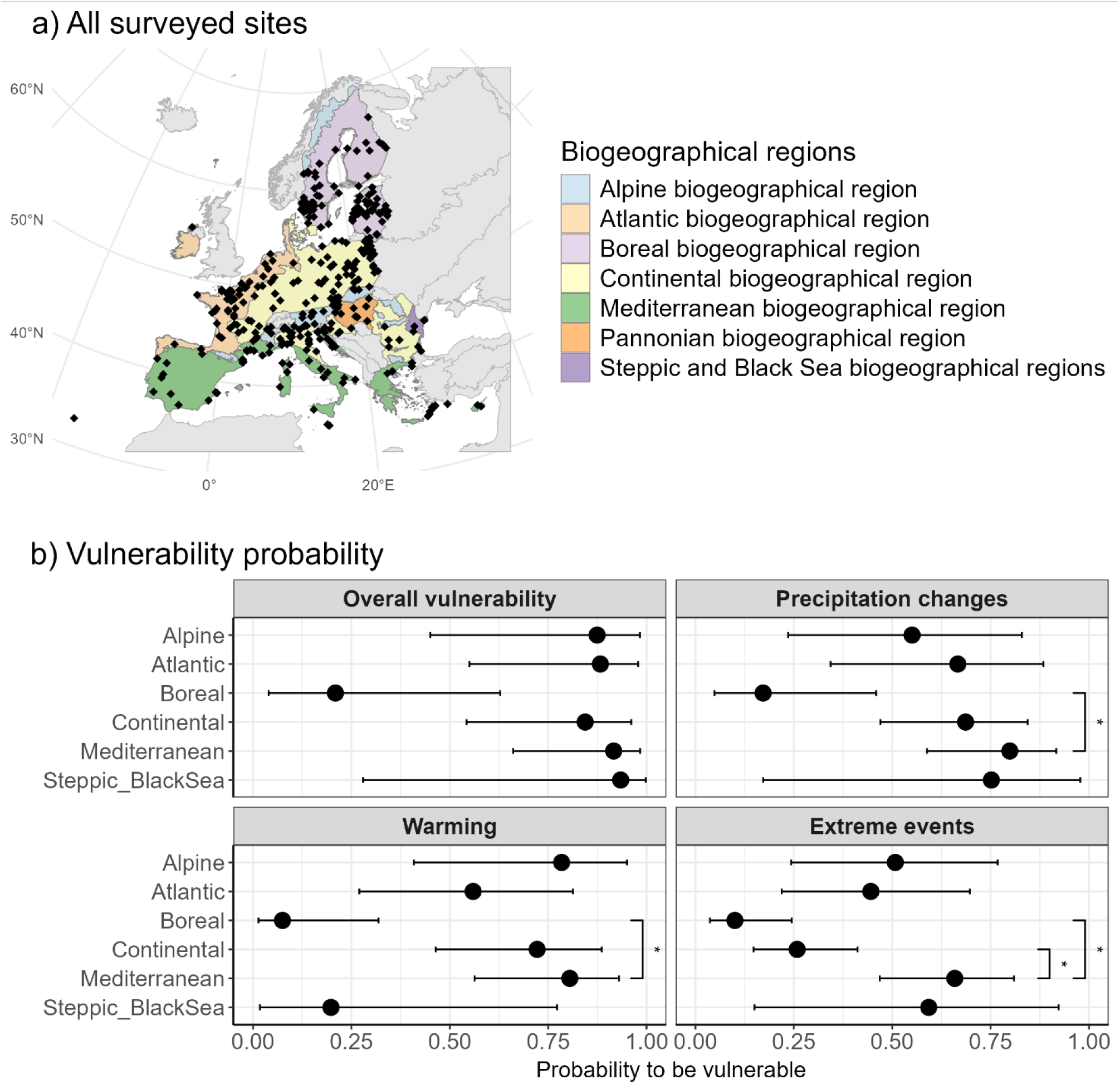
a) Surveyed Natura 2000 sites across European biogeographical regions. One location is shown for multiple overlapping sites. b) Predicted probability (± 95% CI) of reporting vulnerability to different climate-change related threats across biogeographical regions. Significantly different comparisons at *p* < 0.05 are marked with *.

### Climate change management

Climate change was considered in the management by 58.4% (223/382) of the respondents. Managers that perceived their site as vulnerable were 2 times more likely to implement climate adaptation strategies (β = 1.85±0.47, *p* < 0.001; Figure 3). However, 29.8% (68/228) of the managers in vulnerable sites reported no adaptation strategies, or were unsure about their presence. Managers that did not implement any climate-change adaptations were more likely to perceive their site as less vulnerable to it (β = -1.16±0.47, *p* = 0.01; Appendix S8a), whereas no clear distinction was found with managers that were unsure if they had implemented adaptation strategies (β = -0.75±0.41, *p* = 0.07; Appendix S8b).

**Figure 3.**
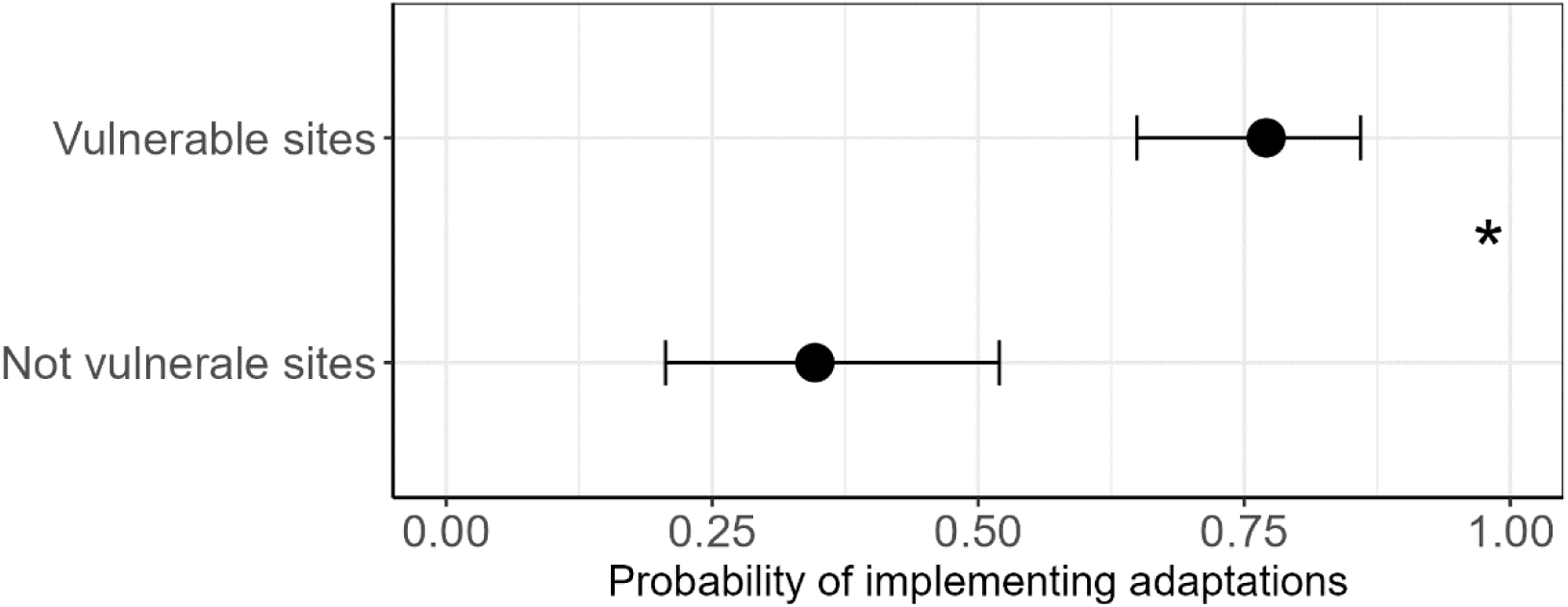
Predicted probabilities (± 95% CI) of implementing climate change adaptation strategies, based on whether the surveyed Natura 2000 site was reported in the previous questions to be vulnerable to climate change or not. Significant difference is marked with * (*p*-value < 0.05).

Conversely to our hypothesis, the implementation of climate change adaptation strategies (done at 223 site) was relatively well balanced between resist, accept and direct approaches (Figure 4). Resisting the effects of climate change (estimated probability = 0.84±0.07) was more frequent than accepting (probability = 0.61±0.11; β= 1.21±0.27, *p* < 0.01), but not more than directing (probability = 0.74±0.1; β = 0.63±0.27, *p* = 0.05). Directing actions were not statistically different than accepting actions (β = 0.59±0.26, *p* = 0.06).

**Figure 4.**
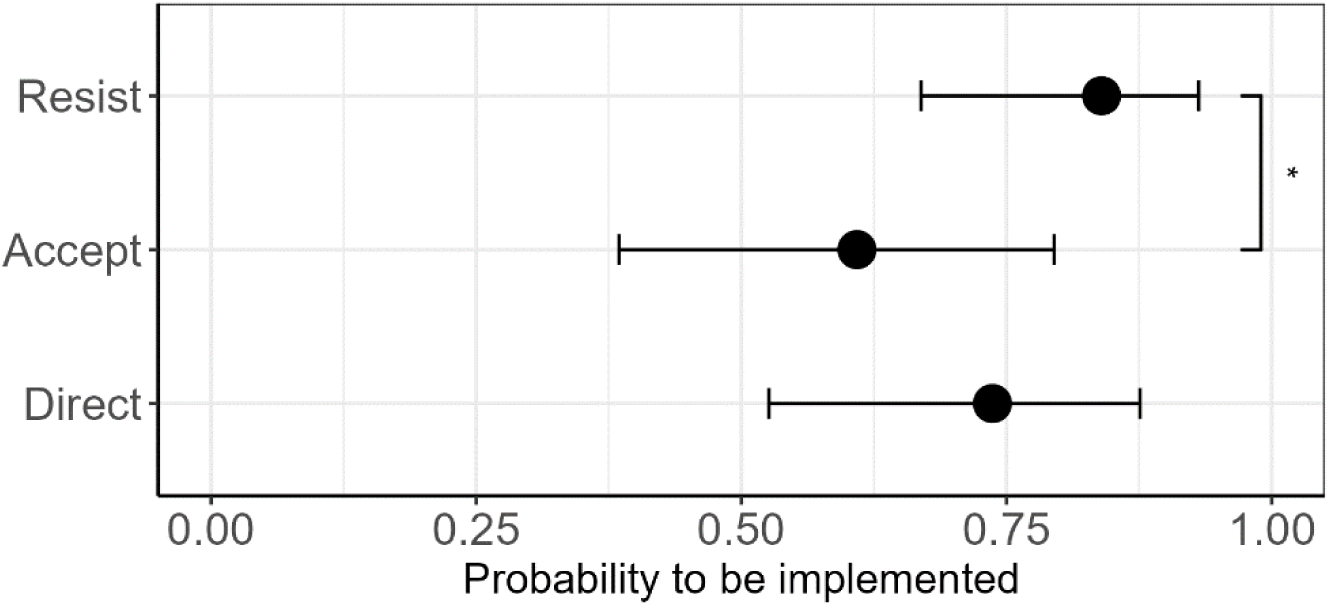
Predicted probabilities (± 95% CI) of the implementation of different RAD approaches (Resist, Accept, Direct) in Natura 2000 sites implementing climate change adaptation strategies (n= 223). Significant differences are marked with * (*p*-value < 0.05).

The most likely reported implementation barrier to climate change adaptation strategies was lack of funding. Lack of knowledge on site vulnerability was more likely to be reported as a barrier than lack of knowledge on RAD actions. The probability of having no issues for the implementation of adaptation strategies was scarce, i.e. 0.08±0.02 (Figure 5; Appendix S9).

**Figure 5.**
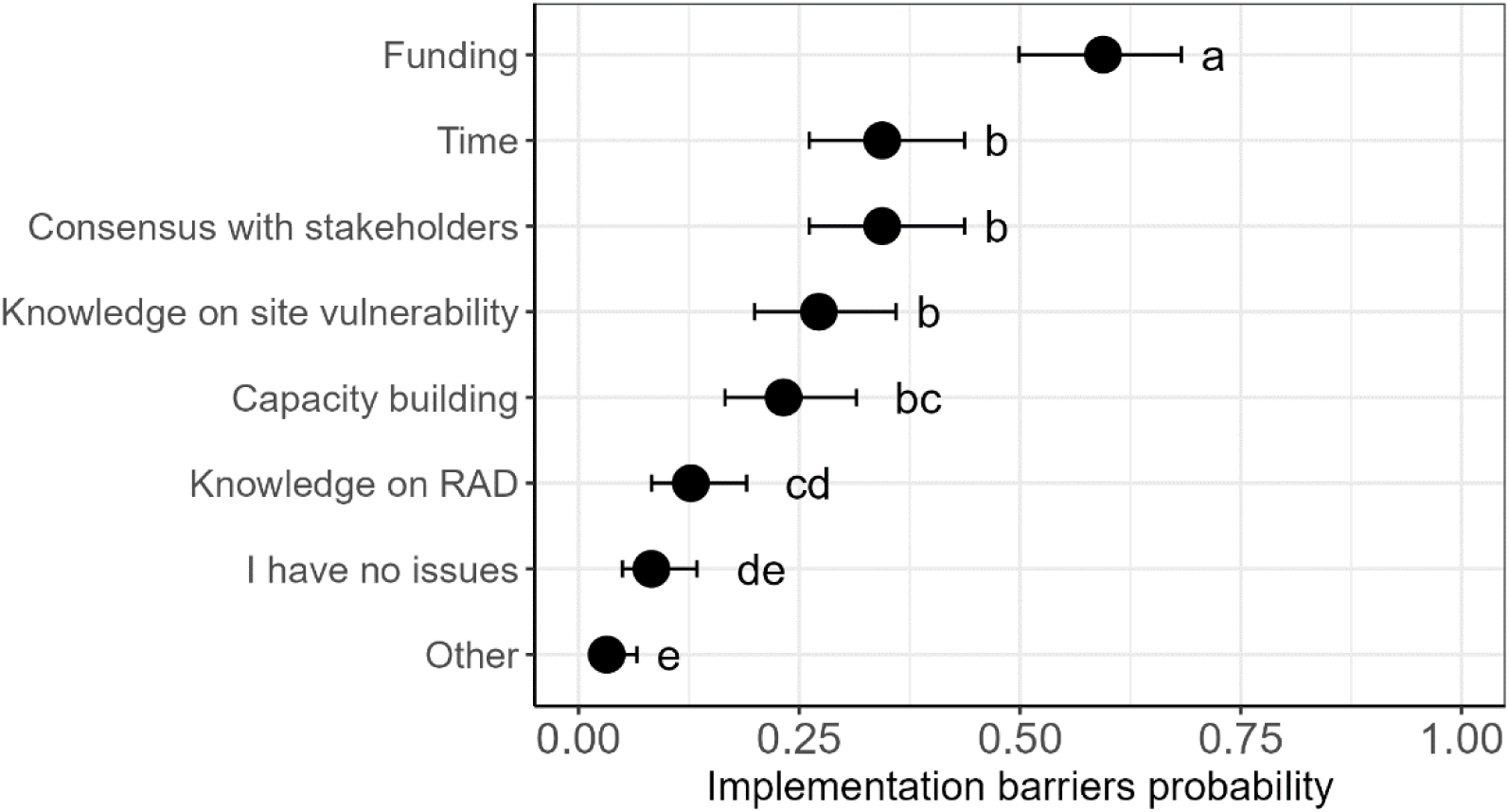
Predicted probabilities (± 95% CI) of different barriers to the implementation of climate change adaptation strategies, considered only for sites that are currently implementing adaptation strategies (n =223). Vulnerability categories whose probabilities are statistically different from each other at *p* < 0.05 are marked with different letters, i.e., shared letters indicate no difference.

## Discussion

We found climate change to be recognised as a frequent threat by N2K protected area managers in the EU, far more than what is reported in the most recent N2K official information for the surveyed sites. Furthermore, vulnerability to climate change varies across biogeographical regions and depends on climate change aspects. Encouragingly, climate change adaptations are already implemented by 58% of the surveyed managers, which is markedly more than what has been found in national studies (Sousa-Silva et al. 2016, EC 2025). Despite reporting funding and knowledge limitations to implement adaptation strategies, we find, contrary to our expectation, that managers use a variety of strategies and not only resist the effects of climate change.

Climate change is a multidimensional phenomenon, so understanding how its various aspects threaten biodiversity on a local scale is essential for effective management (Pearce-Higgins et al. 2019). We found N2K managers’ main concern oriented towards changes in temperature and precipitation rather than extreme climatic events. Warming and precipitation shifts are indeed strong drivers of biodiversity changes in the long-term, with well documented effects (VanDerWal et al. 2013). In comparison, extreme events received less attention in scientific and conservation studies (but see Maxwell et al. 2019), even though they can contribute considerably to biodiversity changes, including on species distribution shifts (Soifer et al. 2025). Extreme events are, however, hard to predict and biodiversity has some potential to recover from them, though limited by the magnitude and frequency of the events and by species-specific characteristics (Sabater et al. 2023; Woodward et al. 2015). While sea level rise was rarely reported as a threat due to its association to coastal areas, it can nonetheless have major impacts on N2K areas (van de Wal et al. 2024; Verniest et al. 2024).

Unexpectedly, in almost all the surveyed N2K sites, managers’ assessment of vulnerability did not correspond to what was reported in the official documentation. With only 7% of consistency between our survey (years 2023-2024) and the latest SDFs update (2021), our results suggest that threat assessments based on the official N2K data could strongly underestimate climate change vulnerability. Similarly, while we expected adaptation strategies to be implemented by a few managers only, 58% of them already did, although a gap persisted since around one third of the vulnerable sites did not implement adaptations. Our survey’s explicit focus on climate change may have biased participation towards managers already engaged with it and the Natur’Adapt LIFE+ campaign promoted the importance of climate adaptations just before our study. While these factors can partly explain the high rates of vulnerability and adaptation strategies, they are unlikely to explain the marked gap between threats reported in our survey and in the SDFs.

### Biogeographical regional contrasts

Managers in the Boreal region were the least worried about each aspect of climate change, while managers in the Mediterranean expressed the most concern. The Mediterranean is a known hotspot of climate change (Tuel and Eltahir 2020), especially due to drought and extreme climatic events (Giorgi and Lionello 2008; Hoerling et al. 2012), and its biodiversity is particularly sensitive to it (Newbold et al. 2020). However, the Boreal region has been warming faster than the rest of Europe (EEA 2009), making our finding of local managers not perceiving it as much vulnerable surprising. Overall, species richness in Northern Europe is expected to increase as a consequence of climate change (Thuiller et al. 2011), though cold-adapted species might be negatively affected (Araújo et al. 2011). Here, management will have the increasingly important role of both accommodating the growing number of warm-dwelling species shifting northwards, while still ensuring that native cold-dwelling species, which have nowhere else to go, still persist.

The survey answers, though well spread across European biogeographical regions (Zavattoni et al., 2025), may still show some spatial clustering, where the same manager may have filled the questionnaire for multiple sites. We corrected the residual spatial autocorrelation in the models to account for potential structuration in the results, though we cannot entirely account for individual perception since we did not collect respondent info. Additionally, even if the answers came from different people, managers working nearby may be influenced by each other’s view or local opinions. Estimating biodiversity vulnerability to climate change is a difficult task that requires in depth knowledge on local climate change exposure and species sensitivity. We provided here an overview of managers’ perceptions, though we could not assess if these align with actual vulnerability, as we would need more details on N2K site conditions.

### RAD implementation

Contrary to our hypothesis, the implementation of resist, accept and direct strategies was relatively well balanced. While strategies based on resisting climate change had a similar probability of implementation as directing, they had a higher probability than accepting. “Resist” has been the traditional, often implicit, approach to conservation which historically viewed ecosystems as static entities (Schuurman et al. 2020). Furthermore, the design of the N2K network possibly favours a resisting approach, since each site is designated with specific species and/or habitat targets, creating a legal obligation for managers to protect these, hence possibly endorsing resistance to change. “Direct” and “Accept” approaches are however used in over half of the sites addressing climate change, highlighting that N2K site managers are open to dynamic management. Resisting is a crucial strategy, especially for conserving species with limited ranges and reduced possibility to shift. However, directing and accepting are needed to more broadly facilitate species responses to climate change, in particular where long term resistance may be costly and unfeasible (Lynch et al. 2021; Schuurman et al. 2020). Ideally, management decisions should be made case-by-case through network level cooperation and trade-offs between different protected areas. If a site accepts the loss of a species due to the changing climate, it is crucial to ensure that the same species still receives enough protection elsewhere.

In this study, we lack details on the implemented conservation measures and data on their effectiveness required to determine the most successful climate change adaptation strategies. Another possible limitation is that the RAD question may have introduced a new concept to some respondents. Thus, despite the explanation provided, not all managers may have been equally confident about categorizing their actions.

### Removing barriers through funding and scientific knowledge

Our results indicate that many barriers exist preventing N2K managers from better implementing climate change adaptation strategies. The most widespread barriers are generic management challenges, not necessarily linked to climate change only, such as lack of funding and time, and difficulties in getting consensus with stakeholders (Geitzenauer et al. 2017; Kati et al. 2015). However, many managers also frequently highlighted more specific barriers, such as lack of knowledge regarding the site’s vulnerability to climate change, which around one-third of the managers reported. Earlier research into the perceptions of N2K forest managers revealed that they found climate change predictions too vague to be useful at a local level and they struggled due to the absence of knowledge on actionable measures (de Koning et al. 2014). These now decade-old findings are echoed in ours and highlight the need to make scientific outcomes more accessible to better support managers’ decision-making processes.

## Conclusions

While reducing greenhouse gas emissions must be a priority, current trends suggest that carbon neutrality will not be achieved soon, making it essential to manage climate change impacts on biodiversity locally. Among the surveyed N2K managers, climate change is already perceived as a widespread threat, which is being managed not only by resisting its impacts, but also by accepting and directing them. However, we identified several knowledge and resource-based barriers in implementing adaptation strategies that must be addressed. Science needs to take an increasingly important role in aiding managers by producing actionable knowledge, while funding and capacity building support are needed to improve research-management synergies and the implementation of adaptation strategies.

## Acknowledgments

We thank the translators, Eurosite, and the managers. The research was funded by the Kone Foundation (Application 202103360) and Biodiversa+ (2021–2022 BiodivProtect program), with the following: Ministry of Environment of Finland (VN/7162/2023), Swedish Research Council (Swedish University of Agricultural Science: 2022-01752), Research Council of Norway (Norwegian Institute for Nature Research: 342927), Innovation Fund Denmark (Aarhus University: 1159-00033B), Swiss National Science Foundation (Swiss Ornithological Institute: 20BD21_209665), and Ministerio de Ciencia e Innovación; Agencia Estatal de Investigación (CREAF: PCI2022-135056-2).

## Data availability statement

The data will be made available on Zenodo upon publication.

