## Supplements for "Vulnerability and adaptations to climate change in EU protected areas: a Natura 2000 managers’ perspective"

#### Appendix S1.

To gather site-level information on how N2K managers perceive and address climate change, we built a survey in R.4.2.1 (R Core Team, 2022) with the ‘shiny’ package and hosted it on Shinyapps (Chang et al., 2022). The survey was translated into European languages (except Romanian, Danish, and Estonian, for which only the English version was available) and active between 1st November 2023 and 15th March 2024. We advertised the survey through various channels, including a partnership with Eurosite (an NGO that connects N2K managers, https://www.eurosite.org/), emails to contacts listed in the Standard Data Form (SDF) European Environment Agency (EEA), and online searches. To encourage participation, we did not request managers to identify themselves, but we asked them to select the name of the N2K site they manage. Since one N2K site can be managed by multiple people, the respondents’ identities could not be fully inferred.

**Appendix S2.** Screenshots of the online survey for Natura 2000 site managers.

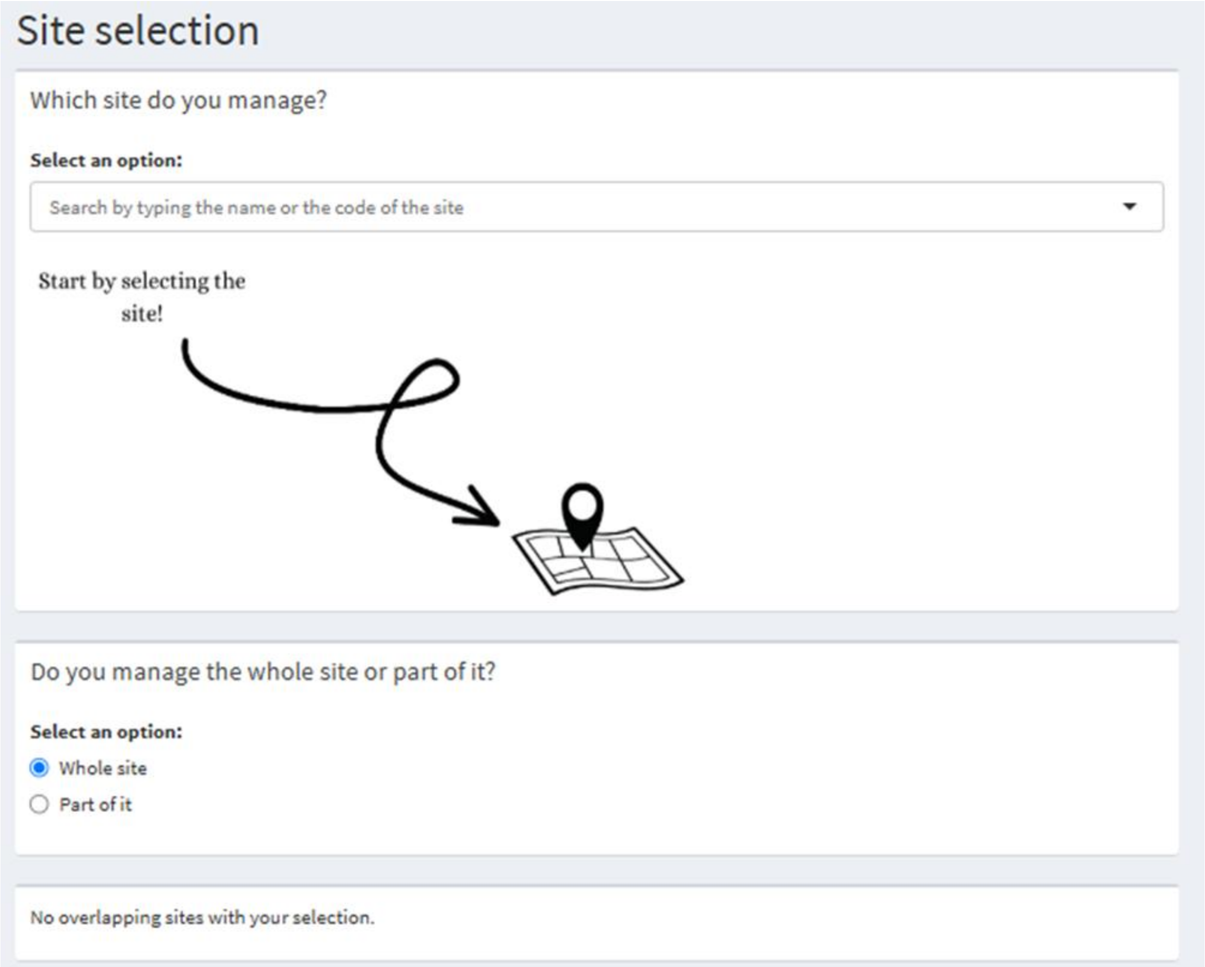
Site selection
Which site do you manage?
Select an option:
Search by typing the name or the code of the site
Start by selecting the site!
Do you manage the whole site or part of it?
Select an option:
Whole site
Part of it
No overlapping sites with your selection.

### (1) Management actions

**Have management actions been done in the site over the last 5 years?**

- (●) Yes
- ( ) No

Here you find the complete list of management actions defined by the European Environment Agency (EEA) divided by category. Please select all the ones you know that have been carried out over the last 5 years (since 2017). For each category you can select multiple actions.

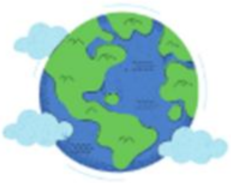

Here we ask you which management actions have been conducted in the site

**Measures related to agriculture and agriculture-related habitats (CA):**

Nothing selected

**Measures related to forestry and forest-related habitats (CB):**

Nothing selected

**Measures related to resources extraction and energy production (CC):**

Nothing selected

**Measures related to development and operation of transport systems (CE):**

Nothing selected

**Measures related to residential, commercial, industrial and recreational infrastructures, operations and activities (CF):**

Nothing selected

**Measures related to the effects of extraction and cultivation of biological living resources (CG):**

Nothing selected

**Measures related to military installations and activities and other specific human activities (CH):**

Nothing selected

**Measures related to alien and problematic native species (CI):**

Nothing selected

**Measures related to mixed source pollution and human-induced changes in hydraulic conditions for several uses (CJ):**

Nothing selected

**Measures related to natural processes, geological events and natural catastrophes (CL):**

Nothing selected

**Measures related to climate change (CN):**

Nothing selected

**Measures related to management of species from the nature directives and other native species (CS):**

Nothing selected

### (1.1) Main management action

Here is the list of management actions you previously selected. In your Natura 2000 site, what is the most important one to protect biodiversity? If there are several actions you consider being equally important, pick one.

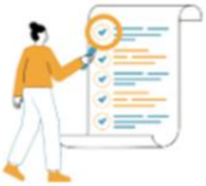

What has been the most important action?

**Select one**

- (●) CA02 Restore small landscape features on agricultural land
- ( ) CA07 Recreate Annex I agricultural habitats
- ( ) CA11 Reduce diffuse pollution to surface or ground waters from agricultural activities
- ( ) CC08 Manage/reduce/eliminate point pollution to surface or ground waters from resource exploitation and energy production

42

Since the year 2000, what is the timing of the action?

**Select:**

- ( ) Permanent
- ( ) Temporary

2000 2001 2023

What percentage of the whole site (including the areas you may not necessary manage) did the conservation action cover (approximatly)?

**Select:**

0 100

0 10 20 30 40 50 60 70 80 90 100

- ( ) Not applicable
- ( ) I do not know

43

#### (1.2) Main management action - threats

Here you find a list of all the threats reported at your site. Which one of these did the management action you previously selected address?

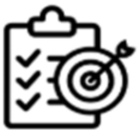

Action ⟶ Threats

**Select:**

- ☐ B04 use of biocides, hormones and chemicals (forestry)
- ☐ A04 grazing
- ☐ G05.06 tree surgery, felling for public safety, removal of roadside trees
- ☐ D01.01 paths, tracks, cycling tracks
- ☐ B02.03 removal of forest undergrowth

Any additional threats addressed by this management action? (leave empty if the answer is no)

**Threat Category:**

Nothing selected

**Specific Threats:**

Nothing selected

#### (1.3) Main management action - species

Here you find a list of the species targeted by the Birds and Habitats Directives reported at your site. Which species are targeted by the management action you previously selected?

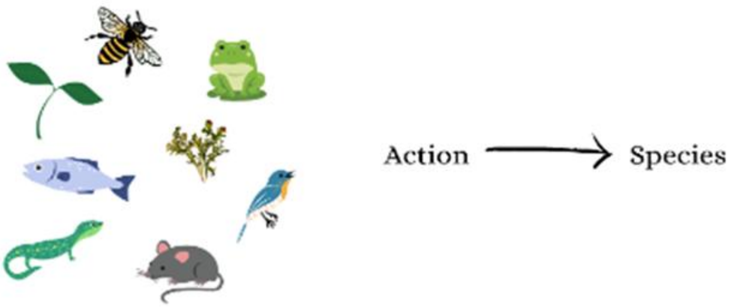

**Bird species**

Select

**Mammal species**

Select

**Reptile species**

Select

**Amphibian species**

Select

45

**Fish species**

Select

**Invertebrate species**

Select

**Plant species**

Select

**Lichen species**

Select

Additional species listed in the Birds or Habitats Directives:

**Please write latin names separated by semicolon (;):**

46

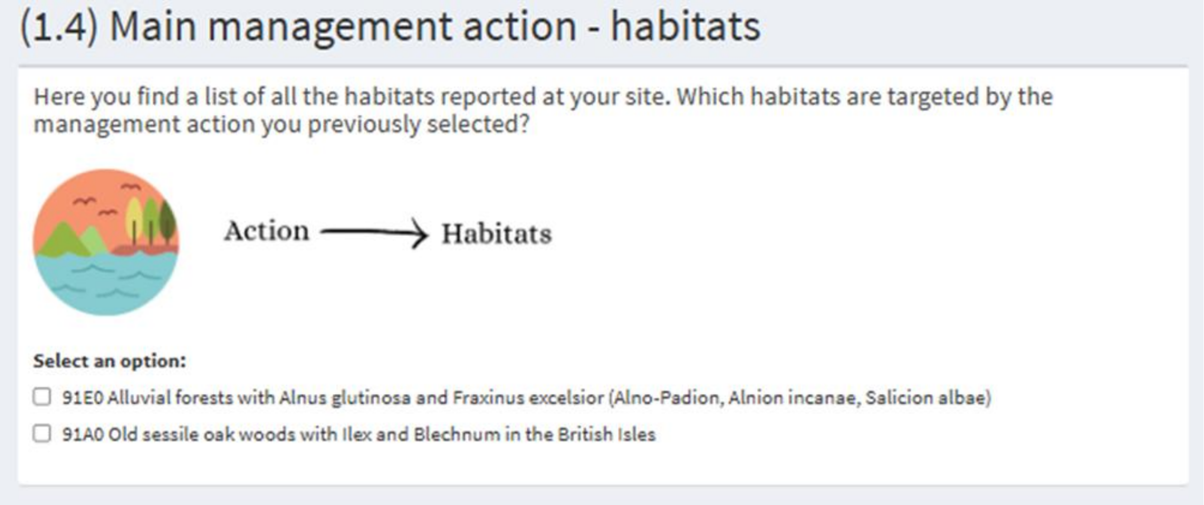

#### (1.4) Main management action - habitats

Here you find a list of all the habitats reported at your site. Which habitats are targeted by the management action you previously selected?

Action ⟶ Habitats

**Select an option:**

- ☐ 91E0 Alluvial forests with Alnus glutinosa and Fraxinus excelsior (Alno-Padion, Alnion incanae, Salicion albae)
- ☐ 91A0 Old sessile oak woods with Ilex and Blechnum in the British Isles

47

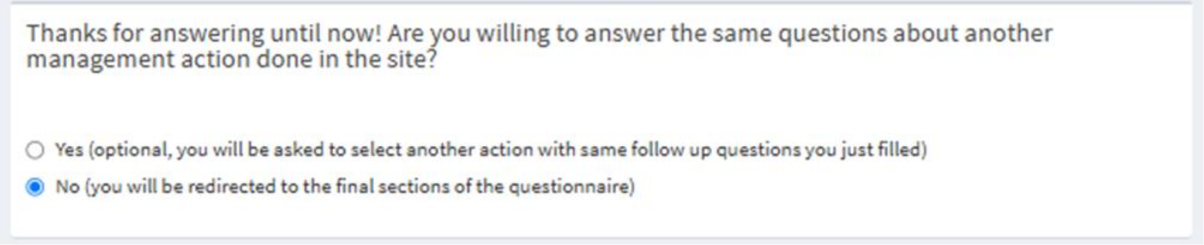

Thanks for answering until now! Are you willing to answer the same questions about another management action done in the site?

- ○ Yes (optional, you will be asked to select another action with same follow up questions you just filled)
- ◉ No (you will be redirected to the final sections of the questionnaire)

48

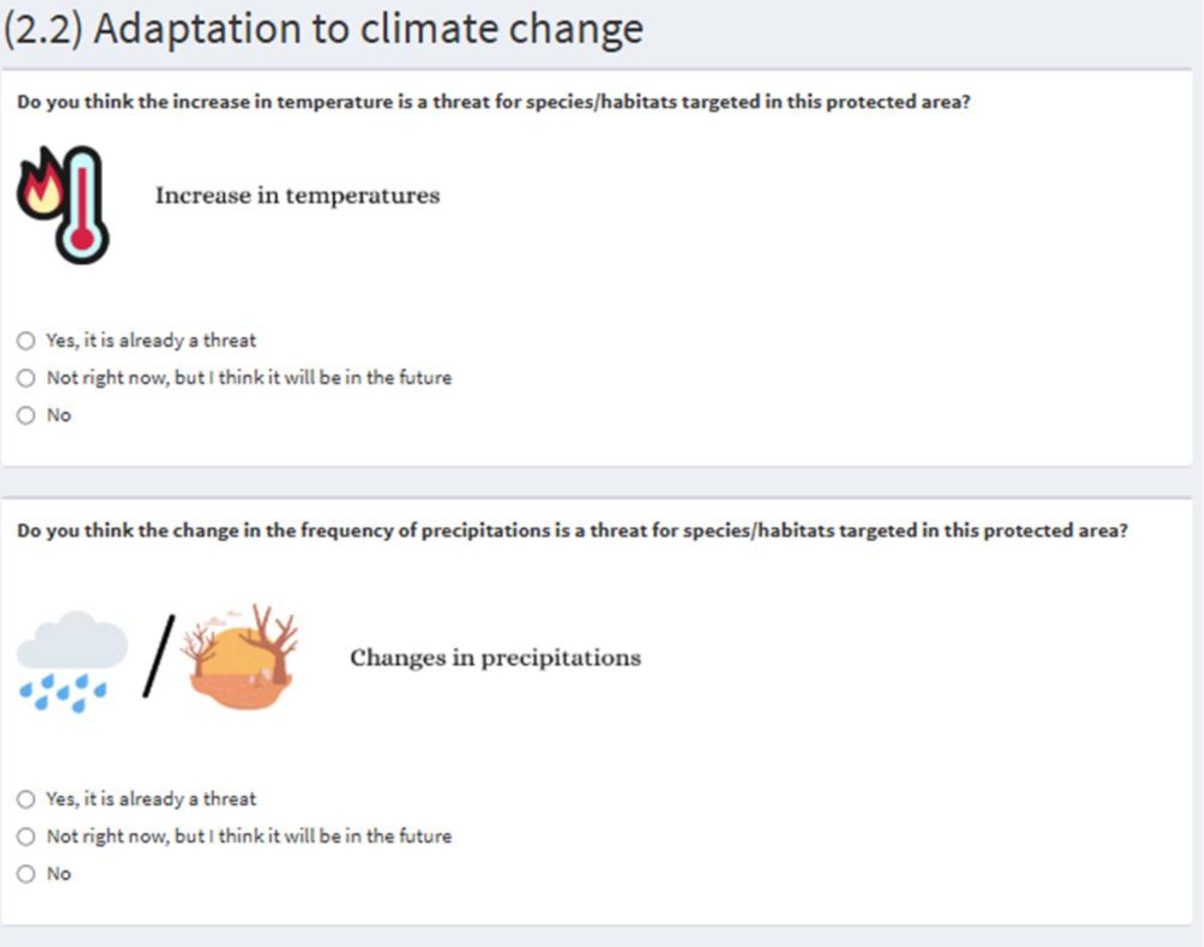

#### (2.2) Adaptation to climate change

**Do you think the increase in temperature is a threat for species/habitats targeted in this protected area?**

Increase in temperatures

- ○ Yes, it is already a threat
- ○ Not right now, but I think it will be in the future
- ○ No

**Do you think the change in the frequency of precipitations is a threat for species/habitats targeted in this protected area?**

Changes in precipitations

- ○ Yes, it is already a threat
- ○ Not right now, but I think it will be in the future
- ○ No

49

**Do you think sea level rise is a threat for species/habitats targeted in this site?**

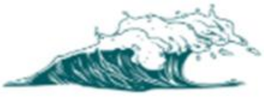

Sea level rise

- ○ Yes, it is already a threat
- ○ Not right now, but I think it will be in the future
- ○ No

**Do you think the increased frequency of climatic extreme events (fires, floods, storms...) is a threat for species/habitats targeted in this protected area?**

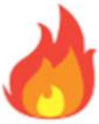

Increase in extreme events

- ○ Yes, it is already a threat
- ○ Not right now, but I think it will be in the future
- ○ No

**Do you consider the impact of climate change in the management of the Natura 2000 site?**

- ○ Yes
- ○ No
- ○ I do not know

Here we ask you about how management actions can help mitigate the effects of climate change in your site following the RAD framework approach (Resist-Accept-Direct, see below). Climate change could be for example temperature increase, precipitation changes, sea level rise or extreme events. The list of possible answers are the actions that you selected before.

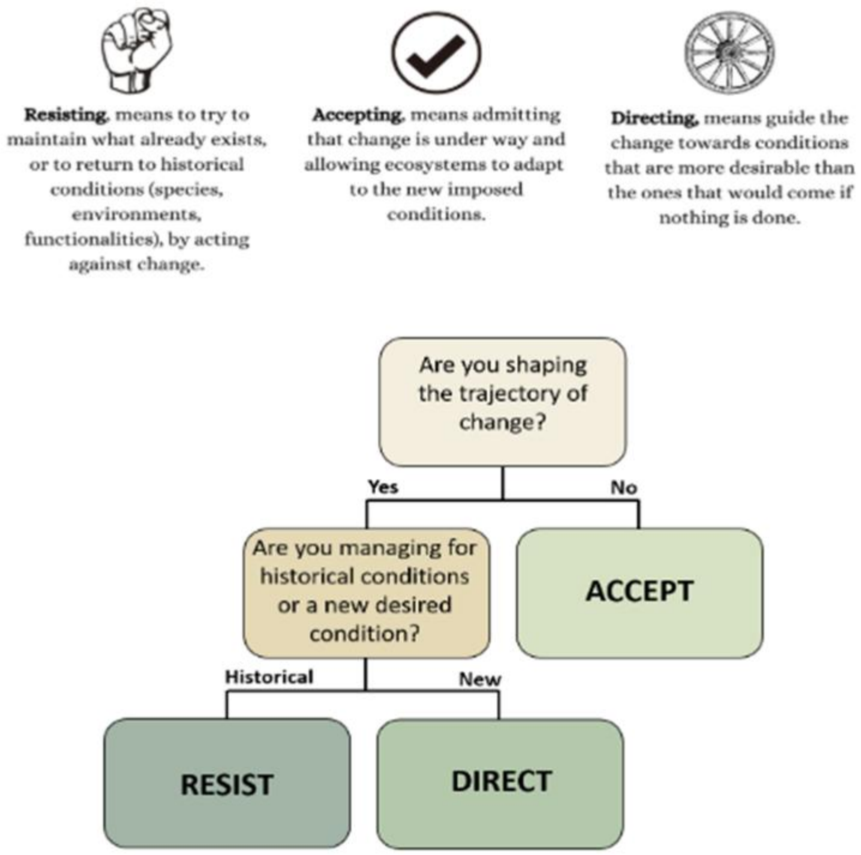

*Images adapted from the LIFE Natur'Adapt and Schuurman et al. (2020).*

☐ The climate change mitigation actions in the site do not fit in this framework

**According to the explanation provided in the picture above, which action(s), if any, have been done in the site to RESIST the effects of climate change?**

☐ There are no resist actions in the site

**According to the explanation provided in the picture above, which action(s), if any, have been done in the site to ACCEPT the effects of climate change?**

☐ There are no accept actions in the site

**According to the explanation provided in the picture above, which action(s), if any, have been done in the site to DIRECT the effects of climate change?**

☐ There are no direct actions in the site

**Is there something preventing you from better implementing actions to deal with the effects of climate change on your site?**

☐ No

☐ Knowledge on site vulnerability to climate change

☐ Knowledge on conservation actions useful to implement the RAD framework

☐ Capacity building

☐ Funding

☐ Time

☐ Consensus with stakeholders

☐ Other

**Is there anything you would like to add about climate change adaptations in the site?**

#### (3) Threats not addressed by management actions

Here you find a list of all the threats reported at your site. Which ones are not addressed by any management actions?

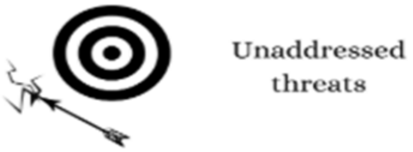

**Select:**

- ☐ B04 use of biocides, hormones and chemicals (forestry)
- ☐ A04 grazing
- ☐ G05.06 tree surgery, felling for public safety, removal of roadside trees
- ☐ D01.01 paths, tracks, cycling tracks
- ☐ B02.03 removal of forest undergrowth

Here you can add additional threats not reported which are present in the site (either addressed by a management action or not):

**Threats related to Agriculture (A)**

Select

**Threats related to Sylviculture, forestry (B)**

Select

**Threats related to Mining, extraction of materials and energy production (C)**

Select

**Threats related to Transportation and service corridors (D)**

Select

53

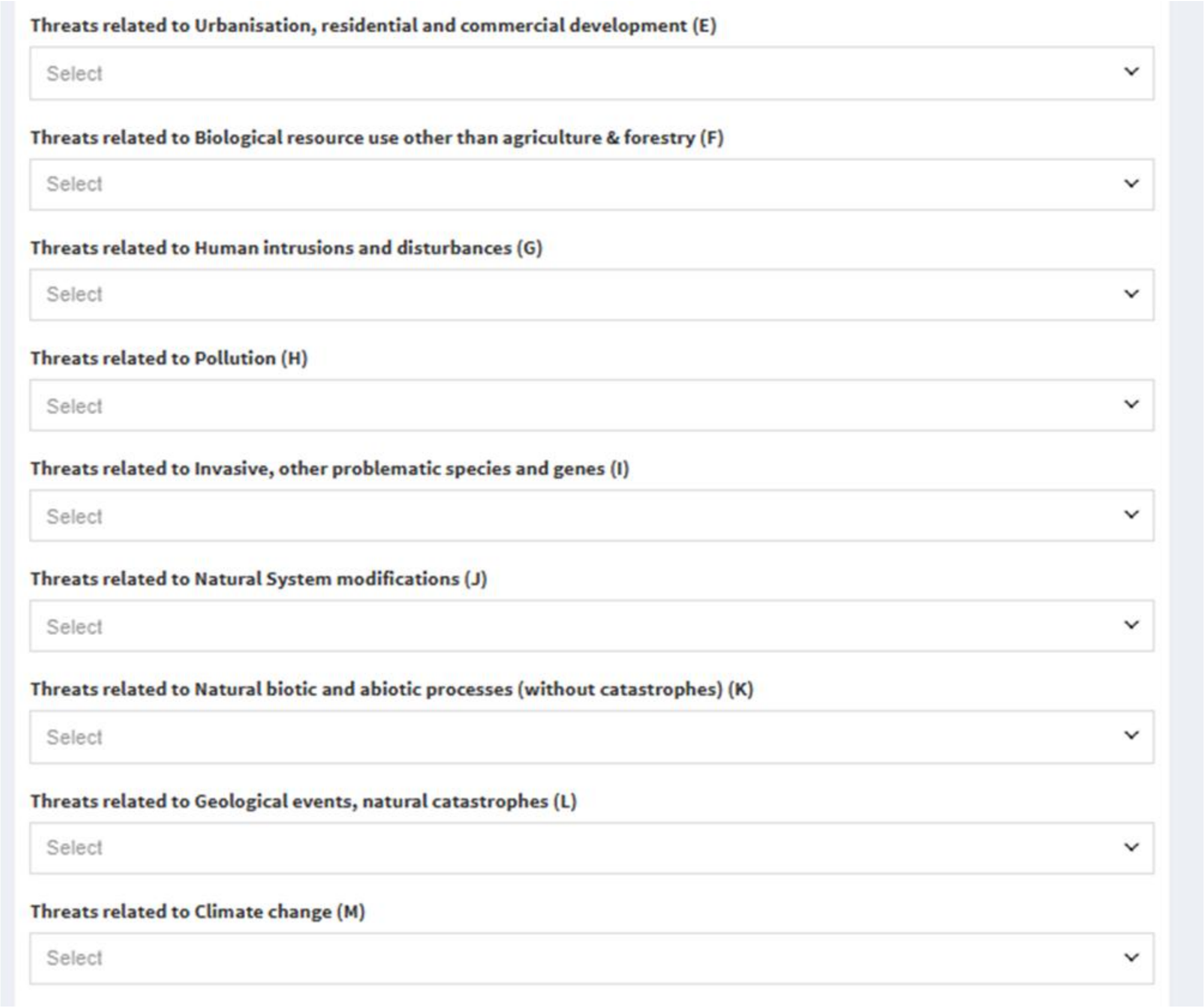
Threats related to Urbanisation, residential and commercial development (E)
Select
Threats related to Biological resource use other than agriculture & forestry (F)
Select
Threats related to Human intrusions and disturbances (G)
Select
Threats related to Pollution (H)
Select
Threats related to Invasive, other problematic species and genes (I)
Select
Threats related to Natural System modifications (J)
Select
Threats related to Natural biotic and abiotic processes (without catastrophes) (K)
Select
Threats related to Geological events, natural catastrophes (L)
Select
Threats related to Climate change (M)
Select

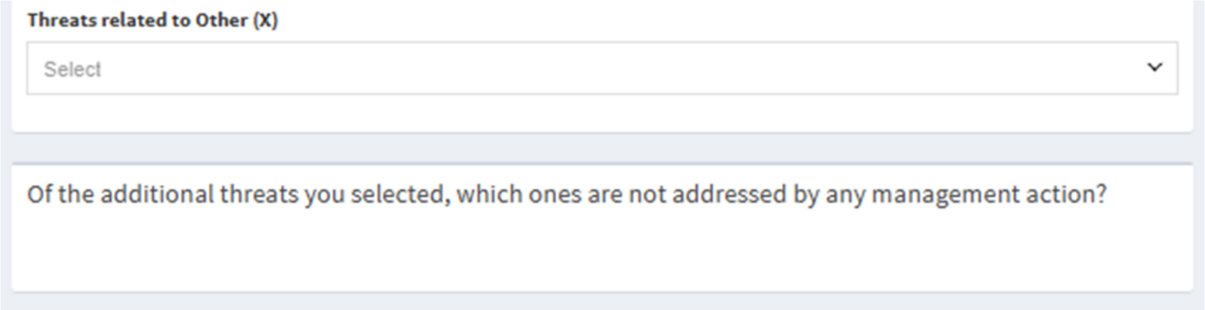
Threats related to Other (X)
Select
Of the additional threats you selected, which ones are not addressed by any management action?

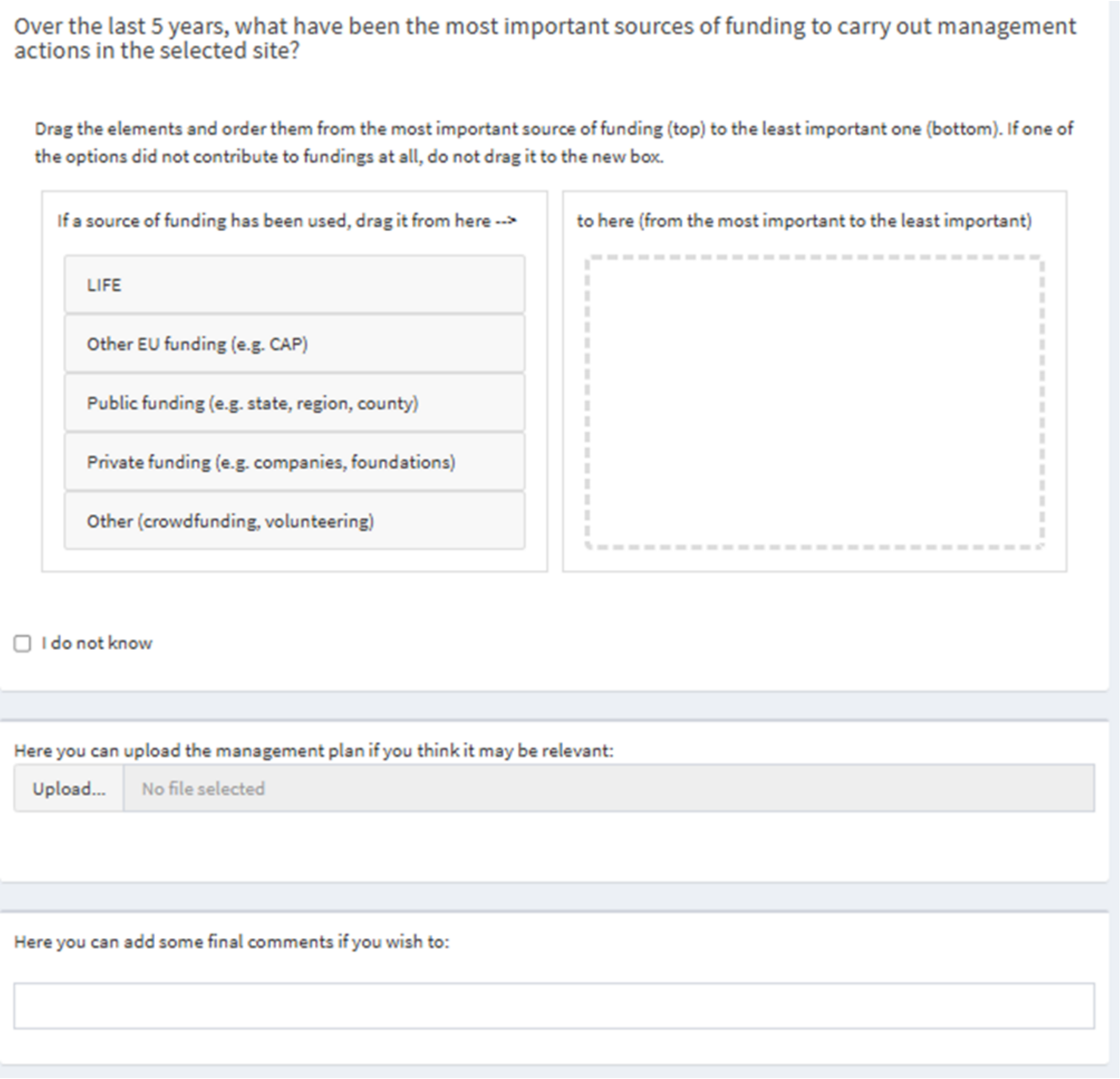
Over the last 5 years, what have been the most important sources of funding to carry out management actions in the selected site?

Drag the elements and order them from the most important source of funding (top) to the least important one (bottom). If one of the options did not contribute to fundings at all, do not drag it to the new box.

If a source of funding has been used, drag it from here -->

LIFE

Other EU funding (e.g. CAP)

Public funding (e.g. state, region, county)

Private funding (e.g. companies, foundations)

Other (crowdfunding, volunteering)

to here (from the most important to the least important)

I do not know

Here you can upload the management plan if you think it may be relevant:

Upload... No file selected

Here you can add some final comments if you wish to:

**Appendix S3.** Comprehensive pairwise comparisons results of the GLM model on climate change – related threats. The response variable is the presence/absence of a threat, while the type of threat is the explanatory variable.

| contrast | estimate | SE | df | t.ratio | p.value |
|---|---|---|---|---|---|
| extreme events – changes in precipitation | -1.470 | 0.235 | 1521 | -6.266 | 0.000 |

| | | | | | |
|---|---|---|---|---|---|
| extreme events – sea level rise | 3.021 | 0.315 | 1521 | 9.582 | 0.000 |
| extreme events - warming | -1.183 | 0.229 | 1521 | -5.154 | 0.000 |
| changes in precipitation - sea level rise | 4.491 | 0.356 | 1521 | 12.599 | 0.000 |
| changes in precipitation - warming | 0.287 | 0.220 | 1521 | 1.310 | 0.557 |
| sea level rise - warming | -4.204 | 0.347 | 1521 | -12.107 | 0.000 |

**Appendix S4.** Comprehensive pairwise comparisons results of the GLM model on overall vulnerability across bioregions.

| **contrast** | **estimate** | **SE** | **df** | **t.ratio** | **p.value** |
|---|---|---|---|---|---|
| Alpine - Atlantic | -0.078 | 1.130 | 366 | -0.069 | 1.000 |
| Alpine - Boreal | 3.268 | 1.732 | 366 | 1.887 | 0.412 |
| Alpine - Continental | 0.252 | 0.863 | 366 | 0.291 | 1.000 |
| Alpine - Mediterranean | -0.454 | 0.937 | 366 | -0.484 | 0.997 |
| Alpine - Steppic_BlackSea | -0.713 | 1.889 | 366 | -0.377 | 0.999 |
| Atlantic - Boreal | 3.345 | 1.548 | 366 | 2.161 | 0.259 |
| Atlantic - Continental | 0.329 | 0.897 | 366 | 0.367 | 0.999 |

| | | | | | |
|---|---|---|---|---|---|
| Atlantic - Mediterranean | -0.376 | 0.968 | 366 | -0.389 | 0.999 |
| Atlantic - Steppic_BlackSea | -0.635 | 1.879 | 366 | -0.338 | 0.999 |
| Boreal - Continental | -3.016 | 1.473 | 366 | -2.048 | 0.317 |
| Boreal - Mediterranean | -3.722 | 1.589 | 366 | -2.342 | 0.180 |
| Boreal - Steppic_BlackSea | -3.981 | 2.251 | 366 | -1.769 | 0.487 |
| Continental - Mediterranean | -0.705 | 0.734 | 366 | -0.961 | 0.930 |
| Continental - Steppic_BlackSea | -0.964 | 1.776 | 366 | -0.543 | 0.994 |
| Mediterranean - Steppic_BlackSea | -0.259 | 1.799 | 366 | -0.144 | 1.000 |

**Appendix S5.** Comprehensive pairwise comparisons results of the GLM model on vulnerability to warming across bioregions.

| **contrast** | **estimate** | **SE** | **df** | **t.ratio** | **p.value** |
|---|---|---|---|---|---|
| Alpine - Atlantic | 1.052 | 1.038 | 366 | 1.014 | 0.913 |
| Alpine - Boreal | 3.804 | 1.422 | 366 | 2.675 | 0.083 |
| Alpine - Continental | 0.336 | 0.820 | 366 | 0.410 | 0.999 |
| Alpine - Mediterranean | -0.127 | 0.876 | 366 | -0.145 | 1.000 |

| | | | | | |
|---|---|---|---|---|---|
| Alpine - Steppic_Blacksea | 2.688 | 1.639 | 366 | 1.640 | 0.573 |
| Atlantic - Boreal | 2.752 | 1.110 | 366 | 2.478 | 0.133 |
| Atlantic - Continental | -0.716 | 0.819 | 366 | -0.875 | 0.952 |
| Atlantic - Mediterranean | -1.180 | 0.849 | 366 | -1.390 | 0.733 |
| Atlantic - Steppic_Blacksea | 1.636 | 1.478 | 366 | 1.107 | 0.878 |
| Boreal - Continental | -3.468 | 1.251 | 366 | -2.772 | 0.064 |
| Boreal - Mediterranean | -3.931 | 1.259 | 366 | -3.124 | 0.024 |
| Boreal - Steppic_Blacksea | -1.116 | 1.511 | 366 | -0.739 | 0.977 |
| Continental - Mediterranean | -0.463 | 0.645 | 366 | -0.718 | 0.980 |
| Continental - Steppic_Blacksea | 2.352 | 1.514 | 366 | 1.553 | 0.630 |
| Mediterranean - Steppic_Blacksea | 2.815 | 1.511 | 366 | 1.863 | 0.427 |

**Appendix S6.** Comprehensive pairwise comparisons results of the GLM model on vulnerability to changes in precipitation across bioregions.

| **contrast** | **estimate** | **SE** | **df** | **t.ratio** | **p.value** |
|---|---|---|---|---|---|

| | | | | | |
|---|---|---|---|---|---|
| Alpine - Atlantic | -0.491 | 0.924 | 366 | -0.532 | 0.995 |
| Alpine - Boreal | 1.774 | 1.079 | 366 | 1.644 | 0.570 |
| Alpine - Continental | -0.581 | 0.707 | 366 | -0.822 | 0.963 |
| Alpine - Mediterranean | -1.176 | 0.786 | 366 | -1.496 | 0.667 |
| Alpine - Steppic_Blacksea | -0.906 | 1.509 | 366 | -0.601 | 0.991 |
| Atlantic - Boreal | 2.265 | 1.092 | 366 | 2.075 | 0.303 |
| Atlantic - Continental | -0.090 | 0.754 | 366 | -0.120 | 1.000 |
| Atlantic - Mediterranean | -0.686 | 0.806 | 366 | -0.851 | 0.958 |
| Atlantic - Steppic_Blacksea | -0.415 | 1.499 | 366 | -0.277 | 1.000 |
| Boreal - Continental | -2.356 | 0.909 | 366 | -2.590 | 0.102 |
| Boreal - Mediterranean | -2.951 | 0.962 | 366 | -3.066 | 0.028 |
| Boreal - Steppic_Blacksea | -2.681 | 1.588 | 366 | -1.688 | 0.540 |
| Continental - Mediterranean | -0.595 | 0.612 | 366 | -0.973 | 0.926 |
| Continental - Steppic_Blacksea | -0.325 | 1.415 | 366 | -0.230 | 1.000 |

| | | | | | |
|---|---|---|---|---|---|
| Mediterranean - Steppic_Blacksea | 0.270 | 1.430 | 366 | 0.189 | 1.000 |

**Appendix S7.** Comprehensive pairwise comparisons results of the GLM model on vulnerability to extreme events across bioregions.

| **contrast** | **estimate** | **SE** | **df** | **t.ratio** | **p.value** |
|---|---|---|---|---|---|
| Alpine - Atlantic | 0.250 | 0.834 | 366 | 0.299 | 1.000 |
| Alpine - Boreal | 2.226 | 0.859 | 366 | 2.592 | 0.102 |
| Alpine - Continental | 1.085 | 0.684 | 366 | 1.587 | 0.608 |
| Alpine - Mediterranean | -0.628 | 0.668 | 366 | -0.940 | 0.936 |
| Alpine - Steppic_Blacksea | -0.344 | 1.207 | 366 | -0.285 | 1.000 |
| Atlantic - Boreal | 1.976 | 0.709 | 366 | 2.788 | 0.062 |
| Atlantic - Continental | 0.836 | 0.612 | 366 | 1.366 | 0.748 |
| Atlantic - Mediterranean | -0.877 | 0.689 | 366 | -1.274 | 0.799 |
| Atlantic - Steppic_Blacksea | -0.594 | 1.216 | 366 | -0.488 | 0.997 |
| Boreal - Continental | -1.140 | 0.579 | 366 | -1.970 | 0.361 |

| | | | | | |
|---|---|---|---|---|---|
| Boreal - Mediterranean | -2.854 | 0.726 | 366 | -3.930 | 0.001 |
| Boreal - Steppic_Blacksea | -2.570 | 1.232 | 366 | -2.086 | 0.297 |
| Continental - Mediterranean | -1.713 | 0.552 | 366 | -3.102 | 0.025 |
| Continental - Steppic_Blacksea | -1.430 | 1.142 | 366 | -1.252 | 0.811 |
| Mediterranean - Steppic_Blacksea | 0.284 | 1.130 | 366 | 0.251 | 1.000 |

**Appendix S8.** Predicted probabilities (± 95% CI) of a) not implementing climate change adaptation strategies and b) being unsure whether climate change adaptations practices were implemented, based on whether the surveyed Natura 2000 site was reported in the previous questions to be vulnerable to climate change or not. Significant differences are marked with * (p-value < 0.05).

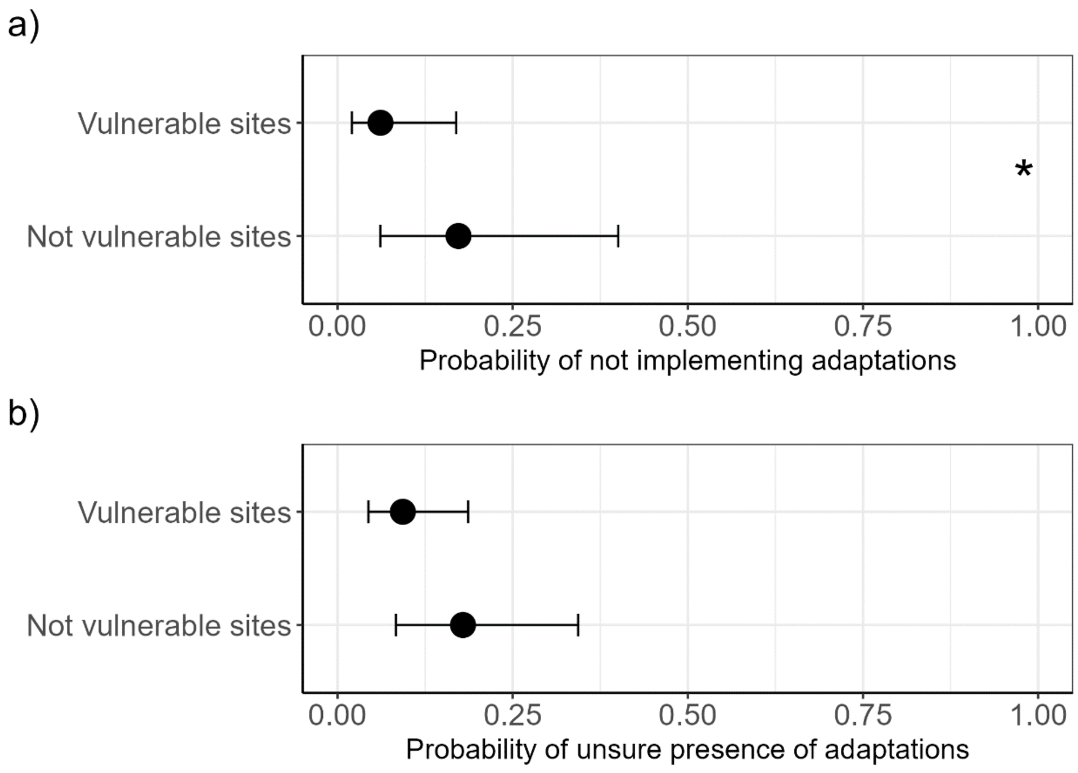

**Appendix S9.** Comprehensive pairwise comparisons results of the GLM model on barriers to climate change adaptation strategies. The response variable is the presence/absence of a barrier, while the type of barrier is the explanatory variable.

| **contrast** | **estimate** | **SE** | **df** | **t.ratio** | **p.value** |
|---|---|---|---|---|---|
| Capacity building - Consensus with stakeholders | -0.550 | 0.226 | 1773 | -2.439 | 0.223 |
| Capacity building - Funding | -1.577 | 0.225 | 1773 | -7.015 | 0.000 |
| Capacity building - I have no issues | 1.212 | 0.293 | 1773 | 4.141 | 0.001 |
| Capacity building - Knowledge on RAD | 0.730 | 0.263 | 1773 | 2.776 | 0.102 |

| | | | | | |
|---|---|---|---|---|---|
| Capacity building - Knowledge on site vulnerability | -0.213 | 0.231 | 1773 | -0.921 | 0.984 |
| Capacity building - Other | 2.218 | 0.401 | 1773 | 5.531 | 0.000 |
| Capacity building - Time | -0.550 | 0.226 | 1773 | -2.439 | 0.223 |
| Consensus with stakeholders - Funding | -1.027 | 0.212 | 1773 | -4.846 | 0.000 |
| Consensus with stakeholders - I have no issues | 1.763 | 0.286 | 1773 | 6.172 | 0.000 |
| Consensus with stakeholders - Knowledge on RAD | 1.280 | 0.255 | 1773 | 5.029 | 0.000 |
| Consensus with stakeholders - Knowledge on site vulnerability | 0.338 | 0.220 | 1773 | 1.533 | 0.790 |
| Consensus with stakeholders - Other | 2.768 | 0.396 | 1773 | 6.990 | 0.000 |
| Consensus with stakeholders - Time | 0.000 | 0.214 | 1773 | 0.000 | 1.000 |
| Funding - I have no issues | 2.790 | 0.287 | 1773 | 9.726 | 0.000 |
| Funding - Knowledge on RAD | 2.307 | 0.255 | 1773 | 9.037 | 0.000 |
| Funding - Knowledge on site vulnerability | 1.365 | 0.219 | 1773 | 6.233 | 0.000 |
| Funding - Other | 3.795 | 0.397 | 1773 | 9.550 | 0.000 |
| Funding - Time | 1.027 | 0.212 | 1773 | 4.846 | 0.000 |

| | | | | | |
|---|---|---|---|---|---|
| I have no issues - Knowledge on RAD | -0.483 | 0.314 | 1773 | -1.535 | 0.788 |
| I have no issues - Knowledge on site vulnerability | -1.425 | 0.289 | 1773 | -4.926 | 0.000 |
| I have no issues - Other | 1.006 | 0.436 | 1773 | 2.307 | 0.290 |
| I have no issues - Time | -1.763 | 0.286 | 1773 | -6.172 | 0.000 |
| Knowledge on RAD - Knowledge on site vulnerability | -0.943 | 0.259 | 1773 | -3.641 | 0.007 |
| Knowledge on RAD - Other | 1.488 | 0.417 | 1773 | 3.571 | 0.009 |
| Knowledge on RAD - Time | -1.280 | 0.255 | 1773 | -5.029 | 0.000 |
| Knowledge on site vulnerability - Other | 2.431 | 0.399 | 1773 | 6.099 | 0.000 |
| Knowledge on site vulnerability - Time | -0.338 | 0.220 | 1773 | -1.533 | 0.790 |
| Other - Time | -2.768 | 0.396 | 1773 | -6.990 | 0.000 |

84
